# Prior knowledge on context-driven DNA fragmentation probabilities can improve *de novo* genome assembly algorithms

**DOI:** 10.1101/2025.03.16.643540

**Authors:** Patrick Pflughaupt, Aleksandr B. Sahakyan

**Affiliations:** MRC WIMM Centre for Computational Biology, MRC Weatherall Institute of Molecular Medicine, Radcliffe Department of Medicine, University of Oxford, Oxford, OX3 9DS, United Kingdom

**Keywords:** *De novo* genome assembly, DNA breakage, genome fragmentation, genome simulations, contigs

## Abstract

**Background:** *De novo* genome assembly poses challenges when dealing with highly degraded DNA samples or ultrashort sequencing reads. Probabilistic approaches have been offered to enhance the algorithms, though existing methods rely solely on expected k-meric frequencies in the assemblies, neglecting the broader sequence context that strongly influences DNA fragmentation patterns.

**Results:** Here, we present a proof of concept showing that prior knowledge on sequence context-driven DNA breakage propensities, through the dedicated parameterisation of k-mer assigned breakage probabilities, can be utilised to recover DNA assemblies that originate from fragmentation patterns more likely to have happened. Our approach is beneficial even for read lengths below the common **∼** 25 bps threshold of modern *de novo* genome assembly algorithms, and well below the threshold used for ultrashort fragments used in ancient DNA research.

**Conclusions:** This work could lay the groundwork for future enhanced *de novo* genome assembly algorithms, with improved ability to effectively assemble and evaluate ultrashort DNA fragments relevant for cell-free, ancient, and forensic DNA research.

## 1 Introduction

*De novo* genome assembly refers to the process of piecing back together the complete genome from short DNA sequence fragments, when no reference genome from a similar species is used or available [1–3]. This process is inherently complex, particularly when dealing with short sequencing reads or a highly degraded DNA, which is common across research in bacterial and viral genomics [4], forensic testing [5], genetic variation [6], natural decay and fossilisation process leading to DNA fragments preserved to a certain degree [7–11]. Highly fragmented genomes typically discard sequencing reads below a threshold of ∼ 35 bps, thus, potentially losing crucial information for complete *de novo* genome assembly. Ultrashort fragments can exponentially bifurcate the assembly graphs, resulting in many possible solutions to the reconstructed sequence and increased computational costs to process them [3]. Consequently, *de novo* assembly algorithms tend to operate within a window of 25 − 50 bp-long sequencing reads, but require very high coverage, which may not always be possible to attain.

To address this problem, a probabilistic-based approach may identify which possible assembled sequence has the maximum likelihood of being closest to the original genome. Various methods have been developed that assess the probability of an assembly hypothesis after taking into account the sequenced data, useful for filling gaps within assembled contigs, addressing errors in base-calling algorithms, and working with non-uniform coverage [12–18]. However, these methods tend to infer probabilities from k-mer frequencies in sequenced reads, which may not adequately account for the intricate probabilistic dependency of DNA breakage from its wider sequence context. This complexity thereby invites the consideration of a novel approach to the *de novo* genome assembly of ultrashort fragments, deriving from our prior research on the context-dependence of various types of DNA double-strand breaks [19].

Here, we present a proof of concept demonstration that prior estimates of k-mer breakage probabilities, i.e. probabilities summarising the k-meric sequence-context effect on the breakage probability at the centre of the k-mer, can be used to assess the fragmentation patterns of the assembled solutions and score them with regard to the probabilities of such fragmentation combinations occurring. We show this by taking, as an example, the breakage probabilities extracted and validated from a range of sequencing experiments [19] (different from the one generated here), with ultrasonication used for the underlying fragmentation. These core probabilities were shown to be independent from the specific experiment, and are prevalently affected by the type of the fragmentation process (see **Figure S13** in [19]). In this work, we then generate many random sequences, and *in silico* ultrasonicate them breaking those down randomly, but biased by those core probabilities to simulate the real conditions for sequencing fragmentation. For each generated sequence, we next assemble the sequences from such simulated fragments using conventional *de novo* sequence assemblers, at this stage omitting any knowledge on the original sequence. We then score all the assembled solutions with and without our proposed additional scoring of the *de novo* solutions against the compliance to the original fragmentation probabilities, which, again, are largely common across experiments that share the same fragmentation process (ultrasonication in this case, but also one can use data from natural decay, biological processes behind cell-free DNA formation of a certain type etc.). Since, in this proof-of-principle work we simulate DNA fragmentation ourselves *in silico*, we have the benefit of actually knowing the full-length identity of the corresponding original sequences generated. This enables us to also rank the solutions with regard to the ground truth, and test our proposed independent metrics for the additional scorings of the solutions.

Our method, which in the brought example is based on probabilities derived from ultrasonication-induced strand breaks, results in a notable improvement in *de novo* genome assembly qualities from ultrasonication-dependent sequencing runs, as compared to assigning uniform probabilities to each base. We expect the same to hold true with parameterisations from any other type of DNA fragmentation process that we have summarised and deposited before (`DNAfrAIlib` from [19]).

## 2 Methods

### General notes on the performed calculations

The developed workflows and analyses in this study employed the R programming language 4.3.2 (https://www.r-project.org). The resource-demanding computations were performed on a single NVIDIA RTX A6000 GPU with 40 GB random access memory. Figures were created with R base, ggplot2 3.4.4 (https://ggplot2.tidyverse.org), and ggsignif 0.6.4 (https://cran.r-project.org/package=ggsignif). Handling of the datasets was done by using R base, tidyverse 2.0.0 (https://cran.r-project.org/package=tidyverse), and data.table 1.14.8 (https://cran.r-project.org/package=data.table). Processing of genomic sequences was done with the Biostrings 2.68.1 (https://bioconductor.org/packages/Biostrings) and plyranges 1.22 (https://www.bioconductor.org/packages/plyranges) libraries.

### Random sampling of genomic sequences

To sample genomic sequences for reference, we took the latest telomere-to-telomere (T2T) human genome assembly version [20] interfaced with BSgenome.Hsapiens.NCBI.T2T.CHM13v2.0 1.5.0 (https://bioconductor.org/packages/BSgenome.Hsapiens.NCBI.T2T.CHM13v2.0). The random sampling was done with a fixed seed of 1234.

### An implementation of a *de novo* genome assembly algorithm

For the purposes of the proof of concept demonstration, we implemented the core *de novo* assembly algorithm into our own toy programme to further test the probability-biased selection of the assembled contigs. As detailed in the further section, it is based on the de Bruijn graph. The model is implemented natively in C++ and interfaced with the R programming language using the Rcpp 1.0.11 library (https://cran.r-project.org/package=Rcpp). More details on the algorithm can be found in the following GitHub repository: https://github.com/SahakyanLab/GenomeAssemblerdev.

### Breakage probability scoring of the assembled sequences

After assembling various solutions, each original sequencing read was aligned against each assembled sequence through brute force alignment, to find the position of the assumed assembled sequence the read originated/broken from. Where a match was found, the broken genomic position was then expanded into an octamer sequence in the given assembly and was assigned a probability of the breakage happening at the centre of that octamer. The overall breakage score of the whole sequence was obtained through the weighted sum of the count and probability of each underlying octamer site where a breakage happened (BPS - breakage propensity score). These calculations were implemented in C++ and interfaced with the R programming language using the Rcpp 1.0.11 library (https://cran.r-project.org/package=Rcpp). We used the widely adopted kseq.h (https://github.com/attractivechaos/klib) for fast sequence parsing and the phmap.hpp (https://github.com/greg7mdp/gtl) for efficient data storage in a hash map data structure. The k-meric breakage probabilities used in this study were derived from mechanically-induced DNA breakage *via* ultrasonication frequencies, obtained from the analysis of [21].

### Evaluating the assemblies against the ground truth real genome

We assessed the quality of our assemblies independently through the above-described breakage propensity score (BPS). As the actual “eye on truth”, to directly assess the similarities between the assembled genomic sequences and the original sequence, which we know owing to the nature of our simulation in this proof-of-principle study, we used the Levenshtein distance metric from the edlib 1.2.7 library [22] in C++ interfaced with Rcpp.

## 3 Results

### 3.1 Sequence context influences the formation of DNA strand breaks

In our recent work, we developed a method to understand how the DNA sequence context may influence the formation of a strand break [19]. We retrieved over a hundred publicly available datasets from various tissues and cell lines, where DNA strand breaks have been studied and their associated genomic positions were reported. Our method involved aligning the breakpoint locations to the initial point of cleavage and calculating their normalised k-meric frequencies. Then, we assessed the variation between adjacent positions using the root-mean-squared deviation (RMSD) metric, revealing k-meric variation in the region surrounding the breakpoint location. Here, our hypothesis posits that the sequence composition and context significantly influence the DNA breakage process, as evidenced by an RMSD peak at the central breakpoint, hence, indicating inherent compositional biases that gradually decay into background levels further from the DNA breakpoint (**Figure S1**). This decay is critical as it signifies the absence of sequence patterns influencing DNA breakages at distant points, hence, representing the background noise to which the signal is expected to converge at a certain distance from the central breakpoint location. We found that, within a 1 kb window, these sequence patterns can be explained by three decoupled range effects (short-, medium-, and long-range) by fitting up to three separate normal distributions to the RMSD signals (**Figure S1** in [19]). The cumulative outcome of these effects forms the full-range sequence influence, where the peak of each curve defines its contribution to their combined effect (**Figures 2, S3, S5-9** in [19]).

### 3.2 DNA breakage probabilities in sequencing fragmentation

Our work identified that DNA strand breaks caused by ultrasound-driven fragmentation [19, 23], normally used in sequencing runs [24], also undergo the above-mentioned three distinct ranges of sequence influence (as per 95% confidence interval used for the definition of the exact boundaries). The immediate neighbouring 14 bases around the central breakpoint location are the most influential, accounting for 72% of a strand break propensities (**Figure S1**). The medium-range effect, covering 93 bp, has a 12% contribution, while the remaining influences come from a longer sequence range spanning 297 bp (**Figures 2** and **S2** from [19]). Given the significance of the short-range sequence effects in forming a strand break, we analysed all the octamer (8-mers) permutations around the central breakpoint. Octamers were chosen as a similar k-meric sequence context (7-mer sequences) was most significant in DNA point mutation works [25–27], while still computationally feasible enough to process, store and retrieve all the parameters. We compared these with negative control octamers located beyond the long-range effect to eliminate any sequence-driven biases. Interestingly, we show that, despite highly correlated and consistent k-meric intrinsic susceptibility scores within replicates of the same breakage type experiment, a comparison of their exact breakage sites revealed that, on average, only 3% overlap (**Figure S15** in [19]). These results highlight that those underlying k-meric breakage probabilities capture the general, and highly consistent, sequence dependence, devoid of any stochastic, or specific/targeted breakage patterns. The breakage probability for a given k-mer was then calculated using the Bayes theorem, as the following ratio [19]:

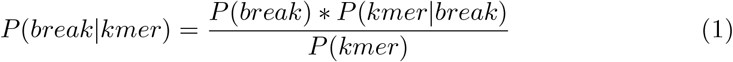

Here, *P* (*kmer*) is estimated through the normalised relative k-meric frequency in the control group, and *P* (*kmer* |*breakage*) through the corresponding frequency in the breakage group.

### 3.3 Simulating context-biased fragmentation of sequences

Ultrasonic frequencies, typically above 20 kHz, are employed for generating shorter DNA fragments necessary for DNA sequencing [21, 23]. In cases where a previously worked-out reference sequence of the same or similar species is unavailable, or when independent assemblies are required for studying genomic rearrangements, fragments are *de novo* assembled into longer-spanning sequences to attempt the reconstruction of the original genome. Although modern algorithms are adept at reconstructing most fragments, some inefficiencies remain, deeming, for instance, the assembly and deposition of a whole genome of a new species, and the full quality assurance still a major milestone in science [28]. Consider a scenario where the *de novo* assembly algorithm outputs two similar sequences but struggles to identify the more accurate one. Here, prior estimates for the sequence-driven probabilities of breakages may reveal which solution would correspond to the DNA fragmentation pattern more likely to have happened, thereby potentially improving the genome assembly quality, and selection of the optimal solution.

As a proof of concept, we simulate the process of fragmenting a genomic sequence into shorter fragments for sequencing, biased by the intrinsic probabilities of breaking between adjacent nucleotides in an ultrasonication setup (**Figure 1**). To do this, we first randomly sampled 200 times 1 kb-long unique DNA sequences across all autosomes from the latest telomere-to-telomere (T2T) human reference genome version [20]. Each of the 200 unique genomic sequences is treated independently. For each sequence, we mapped the breakage probability in an eight-nucleotide sliding window in strides of one nucleotide by using our parameterisation stemming from the `DNAfrAIlib` library (https://github.com/SahakyanLab/DNAfrAIlib), developed as part of our previous study [19]. Hence, by exploiting this library, the probabilities can be retrieved and used for other types of DNA breakage phenomena (natural decay from ancient DNA, forensic DNA, cell-free DNA) behind the DNA fragments to be assembled. To simulate the ultrasonication process, we randomly sampled breakpoint positions biased by their probability of breaking. We repeated this process to achieve a target coverage of ∼ 40 × to avoid sequencing depth being a limiting factor in the *de novo* genome assembly process. Typically, sequencing short fragments follows a process called sequencing by synthesis. To set up the sequencing run, the user specifies the read length, which determines the number of cycles of nucleotide addition. Each cycle adds one nucleotide to the growing DNA strand and snapshots the fluorescence image. To simulate this process, we selected the randomly sampled breakpoint locations and generated reads of a fixed length. Importantly, *de novo* genome assembly algorithms operate within a window of optimal read lengths. Below a threshold, of typically 25 bps, performance quickly diminishes and is often discarded in the quality control step as the assembly algorithms have no natural way to identify neighbouring reads and grow such short reads to a wider sequence context. Similarly, read lengths have a natural ceiling as the sequencing quality decays towards the end of the read. Thus, we simulated this process over varying lengths *l* ∈ (16, 18, 20, 25, 40) with particular emphasis on ultrashort read lengths below the typical lower threshold to demonstrate that the quality of assembled solutions for such extreme cases may improve with our breakage probability-based approach.

**Fig. 1:**
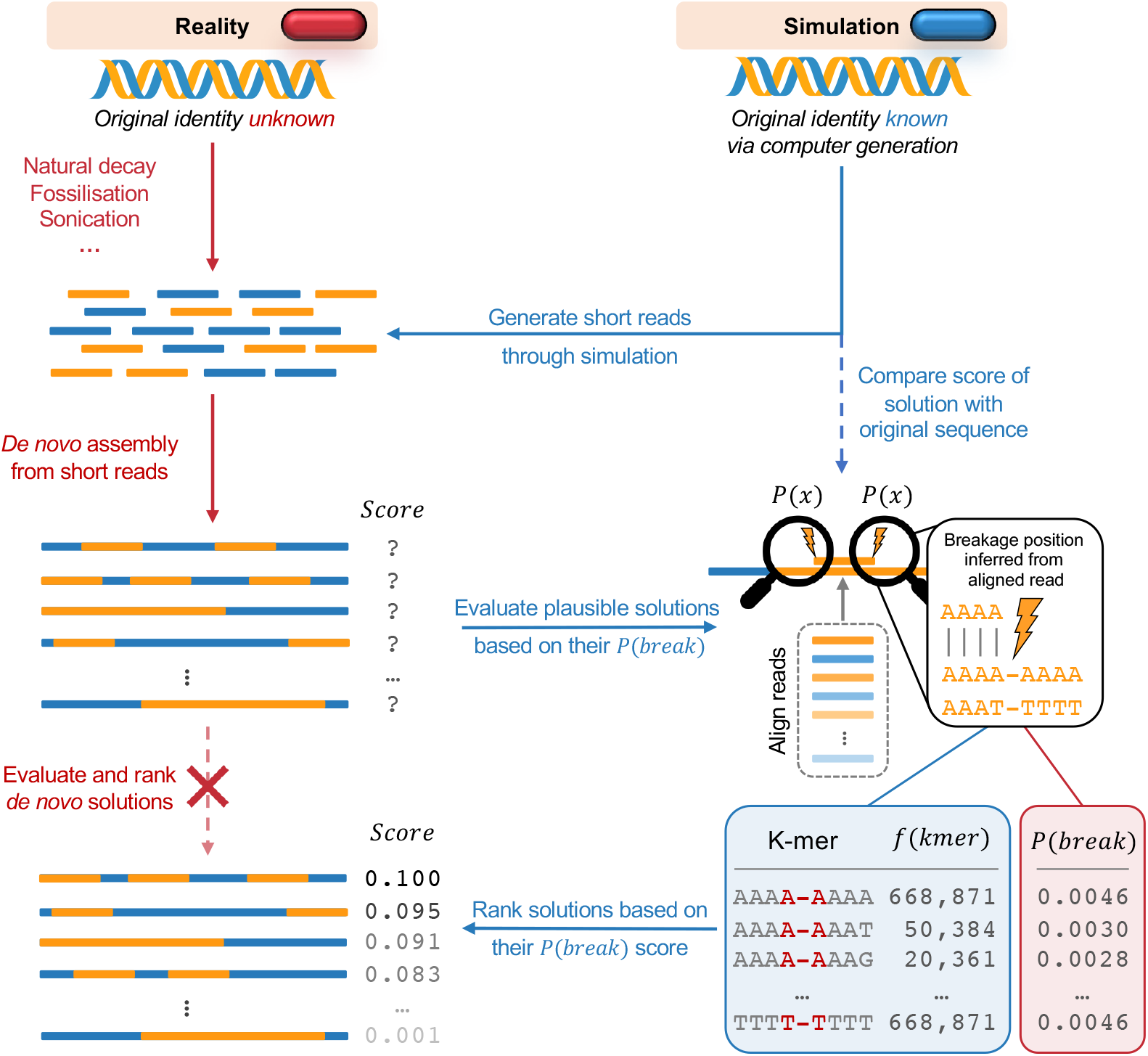
The general depiction of the workflow in this study. The genome is simulated to stochastically break into small fragments biased by the breakage probabilities derived from ultrasonication experiments. This is done to *in silico* (right side, blue pill) simulate the DNA fragmentation process, while actually knowing the real solution for later validation, in contrast to the real world scenario (left side, red pill), where the true sequence is unknown. The generated short fragments are then *de novo* assembled into contigs using our own toy model that is based on the de Bruijn graph. All generated prior sequencing reads are then aligned against each contig solution, to infer the fragmentation pattern which should have been present should those assembled contigs be true. The breakage frequency of each octamer is counted, using the sequences extended around the revealed fragmentation breakpoints. Those frequencies are then contrasted with the octameric breakage probabilities from our prior parameterisation, resulting in a breakage propensity score (BPS) corresponding to the weighted sum of each constituent breakpoint, for each assembly solution, as per its unique originating fragmentation pattern. To evaluate the effectiveness of our breakage scoring metric, each *de novo* assembled solution is then compared against the original genome (known in advance in our simulated scenario) by comparing their sequence similarities *via* the Levenshtein distance metric [22]. A successful proof of concept is expected to have a relationship showing a higher BPS for lower Levenshtein distances. In other words, better contigs that match the reality (smaller Levenshtein distance), have breakage patterns from positions with higher breakage scores, resulting in higher BPS values for breakage patterns.

### 3.4 Selecting for optimal *de novo* assembled sequences

We implemented the core genome assembly algorithm into our own implementation, which generates *de novo* assemblies from our provided sequencing reads. Briefly, it is based on the de Bruijn graph, where a k-mer dictionary is built from the DNA fragments. The de Bruijn graph is a commonly used approach to implementing *de novo* genome assembly algorithms, such as Velvet [29], ABySS [30], and ALLPATHSLG [31]. Hence, for the sake of simplicity and transparency, our proof of concept is a simple implementation of the de Bruijn graph. Traversal of the graph results in long 6 contiguous sequences (contigs). For the simplicity in proof of concept demonstration, we brute-force assembled the contigs into scaffolds, which were then evaluated and ranked based on the underlying sequence context fragmentation patterns (**Figure 1**). While various *de novo* genome assembler algorithms exist, our simple implementation allowed us to isolate the probability-based evaluation of assembled scaffolds from an otherwise lack of criterion in contig assembly.

We next evaluated the breakage probability score (BPS) for each assembled sequence. Here, we aligned each read to the assembled sequence, assigned the octameric breakage probability, and counted the breakage frequency of each octamer. As such, by incorporating a simple scoring metric like the BPS, we can weigh the k-meric breakage counts by the probability of breaking the k-mer, hence, making it one of the most direct, but not necessarily the best, ways to incorporate prior knowledge of k-meric breakage probabilities. At this stage, in practice, the original identity of the assembled sequence is unknown, hence, we would only have access to the BPS metric. However, in this proof of concept demonstration, we can compare the quality of each assembled sequence to the 1 kb-long original sequence (**Figure 1**), which we know due to the nature of our simulation, *via* the Levenshtein distance [22]. Importantly, this comparison is done independently of the *de novo* genome assembly process and serves as a validation for the effectiveness of our breakage scoring methodology in selecting a better assembly solution.

Our results demonstrate that, overall, the majority of the assembled sequences are rather close to the original ones (i.e. the assembly techniques are good for standard conditions), based purely on their breakage compliance to the prior known octameric patterns (**Figure S2**). In particular, we can see that our simulated fragmentation from ultrasonication-induced strand breaks resulted in higher BPS values signifying the compliance of the assemblies with the known sequence biases in ultrasonication-induced fragmentation. This is more visible when, as a control, we compare the BPS distribution with that from solutions stemming from a fully random fragmentation (**Figure S2**), i.e. while defining the breakage positions for the fragmentation using random probabilities drawn from a uniform distribution. This observation is consistent across all read lengths *l* ∈ (16, 18, 20, 25, 40). As expected, this effect becomes more prominent as the sequencing read lengths get shorter in length, as indicated by the increasing statistical significance (**Figure S2**), highlighting the potential to leverage the BPS metric as a criterion for selecting optimal *de novo* genome assembly solutions.

### 3.5 Breakage pattern scoring selects for better *de novo* assemblies

Encouraged by these results, we tried to see whether our breakage pattern scoring can differentiate the optimal assembled sequences from the worse solutions, based only on our k-meric breakage probability compliance through the BPS metric. To that end, we selected the top 5% of sequences based on their BPSs and compared them to the remaining ones. Our results show that the BPS can statistically separate the two groups of assembled sequences (**Figure S3**). Given the variation in sequencing lengths across all assembled solutions, where longer sequences inherently see higher breakage frequencies and thus, BPS, we also normalised the breakage score to its total number of breakpoints (**Figure S4**). Our results show that the differentiation between the top and remaining assembled sequences remains across all read lengths (**Figure S4**).

To demonstrate that BPSs indeed differentiate the better solutions, we binned the Levenshtein distance and evaluated their associated BPS metric (**Figure 2A**). Our results show that our BPS metric tends to increase as the Levenshtein distance decreases, suggesting that the highest quality *de novo* assembled sequences are also most similar to the original genome (**Figure 2A**). Importantly, this trend holds true in sequencing read lengths used in ancient DNA research, and even extends to much shorter reads (**Figure 2A**). While the tendency is consistently strong in (**Figure 2A**), upon normalising the BPS metric by the length of the assembled sequence, the overall trend is still preserved (**Figure 2B**), though with weaker effects in between (sequencing read lengths 18 and 20).

**Fig. 2:**
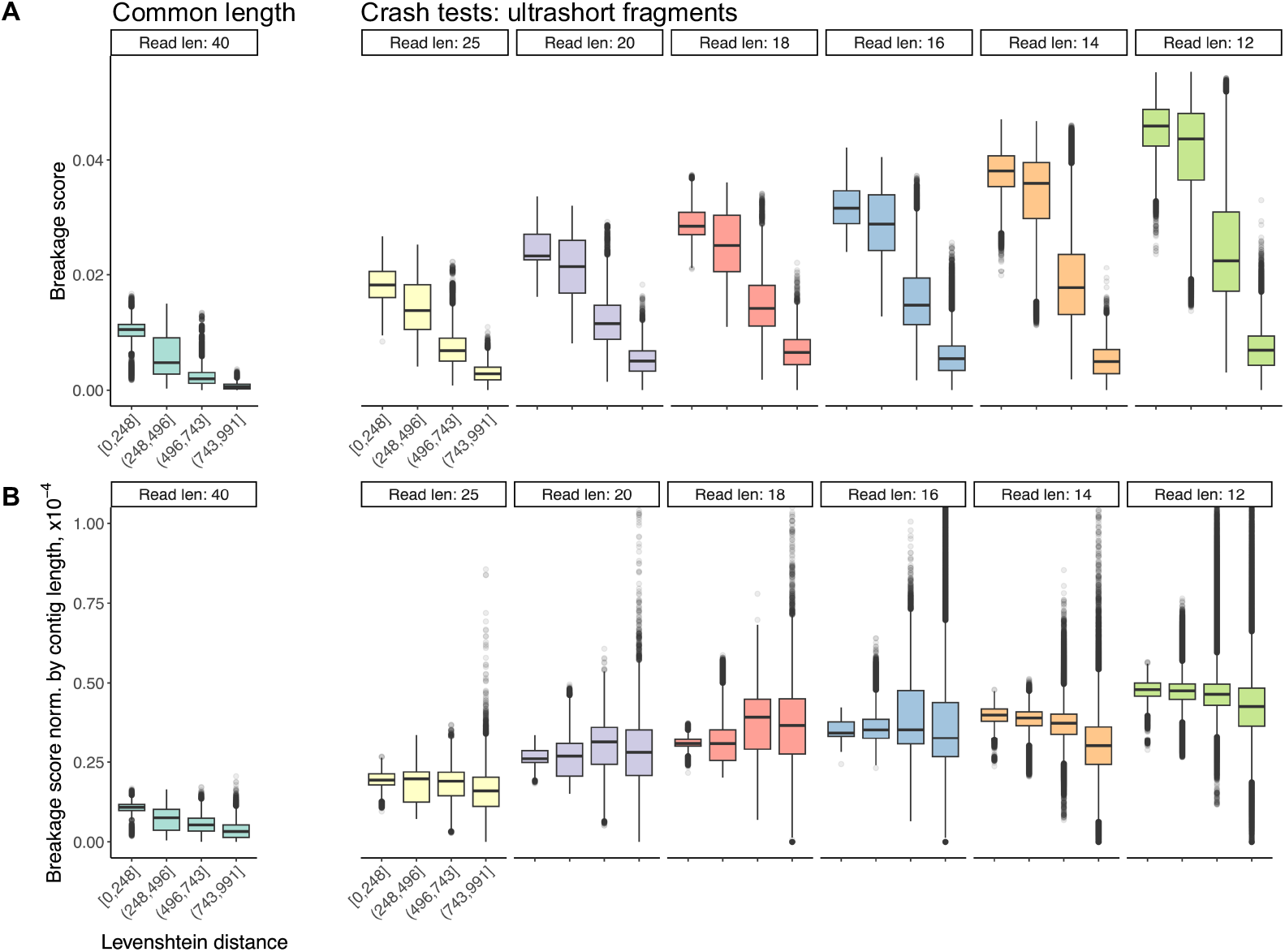
Breakage scores can differentiate optimal *de novo* assembled solutions. The breakage propensity score (BPS) is calculated for each *de novo* assembled sequence, inferring the potential breakage sites from the underlying reads, to assign breakage probabilities necessary for the scoring. To evaluate the effectiveness of our breakage scoring metric, each *de novo* assembled solution is compared against the original genome by comparing their sequence similarities *via* the Levenshtein distance metric; this is binned along the x-axis and presented as boxplots. (**A**) The BPS metric (y-axis) tends to increase with lower Levenshtein distance (x-axis), indicating that the breakage score can select optimal *de novo* assembled solutions. (**B**) The BPS metric is normalised by the length of the assembled solution (y-axis). Generally, the overall trend as in (**A**) is preserved, though with weaker effects in between (read lengths 18 and 20). Overall, the BPS still has the potential to be an effective selection criterion in *de novo* genome assembly problems.

## 4 Discussion

Our results show that the breakage propensity scores (BPSs) tend to increase as the Levenshtein distance decreases, suggesting that the highest quality *de novo* assembled sequences are also most similar to the original genome. Overall, the simplified BPS metric in this work demonstrates that the prior knowledge on fragmentation probabilities can indeed be leveraged as an effective criterion for selecting or for arrival to the optimal *de novo* assembled solutions. As such, integrating the prior estimates of k-meric breakage probabilities directly into *de novo* genome assembly algorithms during the tree traversal process could potentially lower the bound on the read lengths to feasibly assemble ultrashort broken fragments, as seen in the natural decay process leading to ancient DNA fragments. We reiterate that we merely present a proof of concept demonstration that such probability accounting may, in principle, improve the genome assembly process. We hope that our current demonstration is picked up by the community that is specifically devoted to developing fully optimised algorithms for the *de novo* genome assembly of genomes, hence leaving it for others to implement it in the best way possible for application to large-scale datasets and beyond. Considering the overall simplistic and unoptimised implementation of the breakage probabilities in the assembly process, done *via* a mere filtering score for the sake of transparency, we refrain from offering any computational benchmarking at its present proof of principle state. Moreover, considering that the k-meric breakage probabilities are derived from the human genome ultrasonication fragmentation process, which originated from NGS sequencing data, we have not tested our method on long-read sequencing platforms. Ancient mammoths were recently found in North America, believed to be up to 1.2 million years old [32]. Despite the high degradation of the DNA, small amounts of DNA were successfully extracted from just 50 mg of tooth powder. This discovery raises an important question: considering our current demonstration capable of processing read lengths of as short as 14 bps, what is the oldest DNA fragment we could feasibly handle? One study revealed that a 252 bp mitochondrial DNA sequence has an estimated half-life of 521 years, corresponding to a decay rate of 5.50 × 10^*-*6^ per nucleotide [33]. They also suggest that, under the best preservation conditions at −5^*o*^C, all DNA bonds would be broken after a maximum of 6.8 million years, potentially extending back to the era of the earliest hominins [34]. Should such ancient fragments ever be sequenced, it is likely that any remaining DNA would be extremely degraded, with ultrashort fragments making up the majority of them. Thus, integrating prior estimates of k-meric breakage probabilities, from the conditions of natural decay and fossilisation, into *de novo* genome assembly algorithms could potentially lower the current threshold of 25 bp read lengths.

## Abbreviations

bp: base pairs
T2T: telomere-to-telomere
BPS: breakage propensity score

## Acknowledgements

PP is grateful to the UK Medical Research Council (MRC), Hertford College, Clarendon Fund and Radcliffe Department of Medicine for supporting his DPhil studies. The Sahakyan Laboratory has been supported by the UK MRC, MRC Strategic Alliance Funding (MC UU 12025).

## Author contribution

PP and ABS conceived and designed the project, PP performed the research and analyses, PP and ABS wrote the manuscript. ABS supervised the project.

## Availability of code, data and materials

The computer code, necessary to simulate the generation of sequencing reads, *de novo* assembling the fragments, and calculating their breakage scores can be accessed through the following GitHub repository: https://github.com/SahakyanLab/GenomeAssemblerdev.

The computer code, necessary to quantify the sequence context k-meric breakage probabilities is based on our previous work, which can be accessed through the following GitHub repository: https://github.com/SahakyanLab/DNAFragilitydev. The developed `DNAfrAIlib` library of k-meric breakage probabilities and fragility scores is publicly available *via* the https://github.com/SahakyanLab/DNAfrAIlib GitHub repository.

## Declarations

### Ethics approval and consent to participate

Not applicable.

### Consent for publication

Not applicable.

