## Supporting Information for "Prior knowledge on context-driven DNA fragmentation probabilities can improve *de novo* genome assembly algorithms"

### Supplementary Information

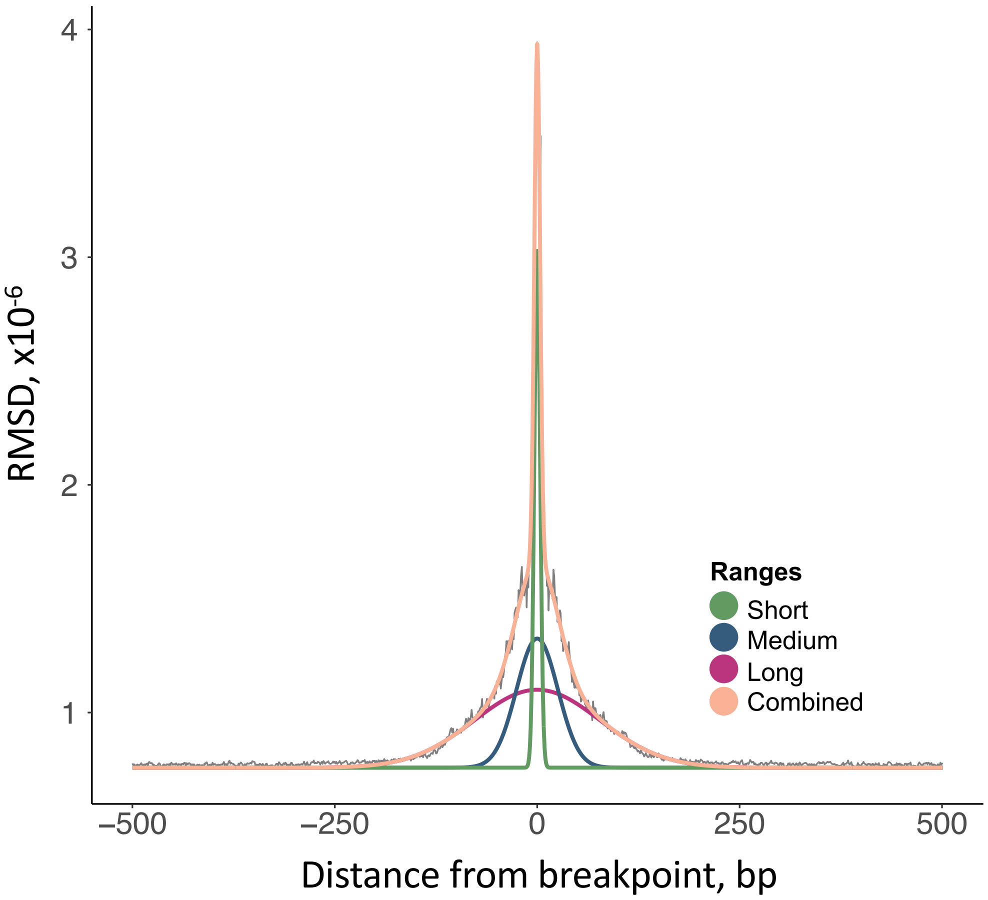


**Fig. S1 Quantifying the span of sequence-based influences for ultrasonication-induced DNA strand breaks.** We calculated the range of sequence-context influence in a 1 kb span surrounding the central breakpoint location generated from ultrasound frequencies. Gaussian curves were fitted to quantify the ranges of influence for each autosome.

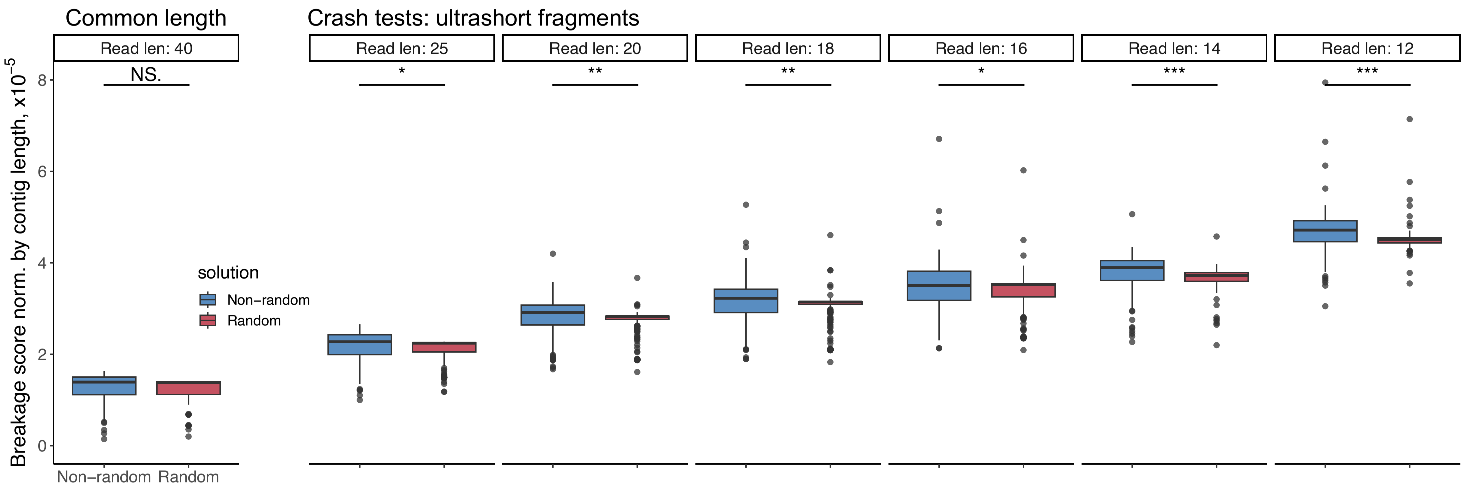


**Fig. S2 The breakage propensity score between the breakage probability distribution of the *de novo* assembled sequences and the reference genome normalised by the assembled sequence length.** We calculated the breakage propensity score for each *de novo* assembled sequences, using the underlying reads to infer the potential breakage positions. The Levenshtein distance between the assembled solutions and the original actual sequence is then compared while using BPSs with actual breakage probabilities, and with randomly sampled breakage probabilities drawn from a uniform distribution (red). The results show that assembling *de novo* sequences using BPSs results in better solutions compared to using random probability values. We performed a two-sample t-test between the two populations, separately for each sequencing read length specified in the experiment. The statistical significance is annotated on top of the bars (******* $p < 0.001$, ****** $p < 0.01$, ***** $p < 0.5$, **N.S.** not significant).

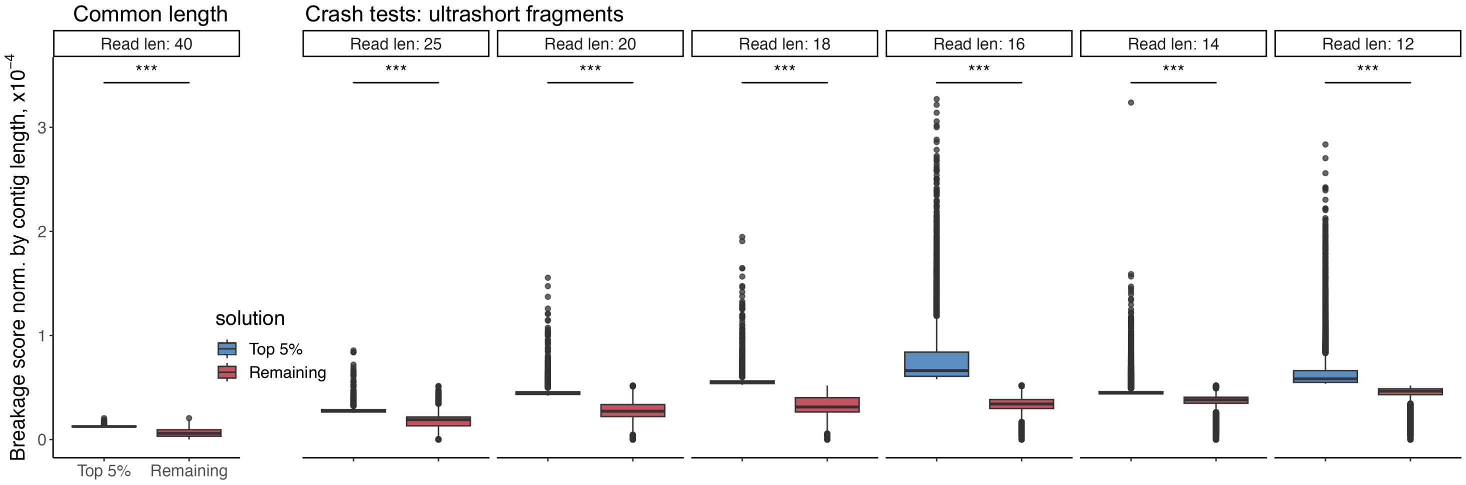


**Fig. S3 The top 5% versus remaining 95% of breakage propensity scores of *de novo* assembled sequences.** The breakage propensity score is calculated for each *de novo* assembled sequence, which is the weighted sum of the k-meric breakage probability and the number of times the k-mer got broken in this *de novo* assembled sequence. The sequences with the top 5% and the remaining 95% are presented in boxplots. We performed a two-sample t-test between the two populations, separately for each sequencing read length specified in the experiment. The statistical significance is annotated on top of the bars (******* $p < 0.001$).

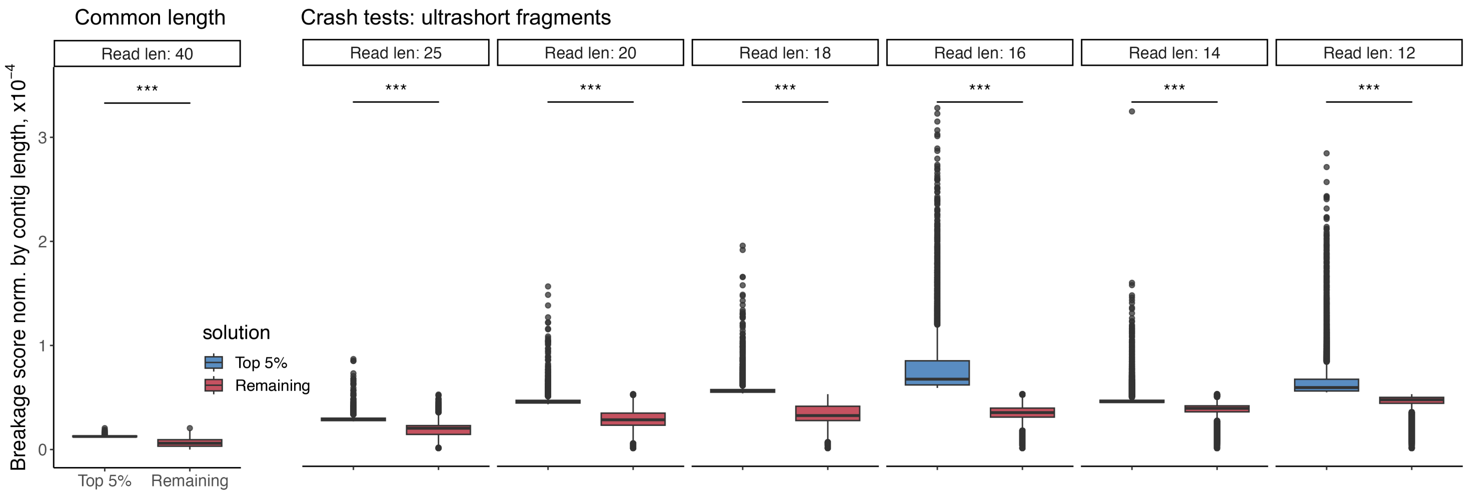


**Fig. S4 The top 5% versus remaining 95% of normalised breakage propensity scores of *de novo* assembled sequences.** The breakage propensity score is calculated for each *de novo* assembled sequence, which is the weighted sum of the k-meric breakage probability and the number of times the k-mer got broken in this *de novo* assembled sequence. This score is then normalised to its total frequency of breaks of the assembled sequence. The sequences with the top 5% and the remaining 95% are presented in boxplots. We performed a two-sample t-test between the two populations, separately for each sequencing read length specified in the experiment. The statistical significance is annotated on top of the bars (******* $p < 0.001$).
